# Role of adenosine in functional recovery following anoxic coma in *Locusta migratoria*

**DOI:** 10.1101/2019.12.19.882464

**Authors:** Rachel A. Van Dusen, Christopher Lanz, R. Meldrum Robertson

**Affiliations:** Department of Biology, Queen’s University, Kingston, ON. Canada

**Keywords:** Spreading depolarization, insect, caffeine, anoxia, motor patterning

## Abstract

When exposed to prolonged anoxia insects enter a reversible coma during which neural and muscular systems temporarily shut down. Nervous system shut down is a result of spreading depolarization throughout neurons and glial cells. Upon return to normoxia, recovery occurs following the restoration of ion gradients. However, there is a delay in the functional recovery of synaptic transmission following membrane repolarization. In mammals, the build-up of extracellular adenosine following spreading depolarization contributes to this delay. Adenosine accumulation is a marker of metabolic stress and it has many downstream effects through the activation of adenosine receptors. Here we demonstrate that adenosine lengthens the time to functional recovery following anoxic coma in locusts. Caffeine, used as an adenosine receptor antagonist, decreased the time to recovery in intact animals and lengthened the time to recovery in semi-intact animals. Our results show that the rate of recovery in insect systems is affected by the presence of adenosine.

## 1. Introduction

The nervous system requires a constant supply of energy in order to maintain proper functioning and thus exposure to metabolic stress can have severe consequences on neural performance. In response to environmental stressors, such as anoxia, some insects are capable of entering a reversible coma via complete neural shutdown. The onset of the coma is associated with the loss of ion gradients and a large redistribution of ions between neurons, glia and the interstitium, a phenomenon known as spreading depolarization (SD). SD has been described in mammals for many years (Leao, 1944) and was more recently identified in insects (Rodgers et al., 2010; Spong et al., 2016). Key characteristics of SD have been well established and are consistent in both mammalian and insect models, however there is still uncertainty about the underlying mechanisms. Recovery following coma depends on the re-establishment of ion gradients across cellular membranes. Several factors can influence the rate of recovery and accumulation of extracellular adenosine contributes to a delay in recovery following SD in mammalian models (Lindquist and Shuttleworth, 2012; 2017). Here we investigated the role of adenosine on the recovery from anoxic coma in an insect model of SD, the locust, *Locusta migratoria*.

Adenosine is an endogenous metabolite that is essential to cellular activity and that influences cellular energy requirements. It can act to protect cells against metabolic stress (Borea et al, 2018). However, in neural tissue extracellular adenosine accumulation results in delayed recovery of synaptic transmission following SD (Lindquist and Shuttleworth, 2012; 2017). Such a delay in recovery of synaptic transmission can lead to increased cellular damage and impaired function. This action of adenosine has been established in mammalian preparations and it has not yet been explored in invertebrate models. Invertebrate adenosine receptors (ARs) have been identified but their amino acid sequences have not been closely conserved and are dissimilar to those of mammalian receptors, with less than 30% amino acid identity (Kalinowski et al, 2003; Dolezelova et al, 2007). Despite sequence variation, functional continuity has been found. Adenosine receptor agonists and antagonists affect invertebrates similarly to mammals (Andretic et al, 2008; Hendricks et al, 2000).

In mammals, the effects of adenosine are mediated via four G-protein receptors, each with varying distribution and affinity for adenosine. High affinity receptors (A_1_, A_2a_, A_3_) are more readily bound with lower adenosine concentrations, whereas the low affinity receptor (A_2b_) is favoured with high concentrations (Fredholm et al, 2001). The A_1_ receptor is the most widely distributed receptor throughout the mammalian central nervous system. Activation of A_1_ results in a depression of excitatory transmission, and also leads to the reduction of cyclic-AMP (cAMP) through the inhibition of adenylyl cyclase (Borea et al, 2018). cAMP activates protein kinase A (PKA) which has known protective effects on CNS function in locusts during thermal stress (Armstrong et al, 2006). Although the signalling pathways following adenosine receptor (AR) activation are largely unknown in invertebrates, the inhibition of adenylyl cyclase may be involved.

Locusts are resilient animals, with the ability to tolerate extended periods of anoxia and recover from comas and SD (Rodgers et al, 2007). This invertebrate model facilitates study of the characteristics of SD at the molecular, cellular, and systemic levels. Anoxic comas can be induced via water immersion, gas, or chemical anoxia (Rodgers et al, 2007). Coma onset is characterized by cessation of patterned neural activity, a ∼40-50 mV drop in trans-perineurial membrane potential (TPP), and a surge of extracellular potassium in the interstitium (Rodgers et al, 2009). During recovery, neural excitability and patterned activity return, along with the reestablishment of membrane potentials. The sequence and rate of recovery are affected by the mode of anoxia and the conditions of the experiment. The return of motor patterns is indicative of synaptic transmission recovery and was measured to observe the effect of pharmacological manipulation on recovery rates.

We investigated the role of adenosine on induction to, and recovery from, anoxic coma via water immersion and nitrogen gas in locusts. Based on mammalian findings, we predicted that AR activation would delay recovery of neural activity, and that AR inhibition would decrease recovery time.

## 2. Methods

### 2.1 Animals

All experiments were performed on gregarious African migratory locusts, *Locusta migratoria migratorioides*, aged 3-5 weeks past the final molt. Animals were taken from a crowded colony located in the Animal Care Facility of the Biosciences Complex at Queen’s University (Kingston, Ontario, Canada). The colony is subject to a 12:12 hr light:dark photoperiod schedule at a room temperature of 30 ± 1 °C during light hours and 26 ± 1 °C during dark hours, and a constant humidity of 23 ± 1 %. Animals were fed wheat seedlings and a dry mixture of 1-part skim milk powder, 1-part torula yeast, and 13-parts bran.

### 2.2 Whole animal water immersion

Male locusts were removed from the colony and submerged in a tank filled with room temperature de-chlorinated tap water. Animals were held in individual compartments and were left submerged for 20 minutes before being taken out of the tank, dried, and weighed.

### 2.3 Whole animal gas anoxia

Appendages were removed, an EMG electrode was inserted into the 3^rd^ abdominal segment 1-2 mm above the spiracle and a silver ground wire was inserted into the anterior portion of the thorax; both wires were secured with small drops of wax. Animals were placed individually into a 50 mL syringe, fitted with a gas exchange at one end and an opening at the other to allow for flow-through (Fig 1). A 5-minute baseline recording was taken with air flowing through the syringe. After the baseline recording, nitrogen gas was turned on for 30 minutes. After 30 minutes the nitrogen gas was switched back to air to allow for recovery.

**Figure 1.**
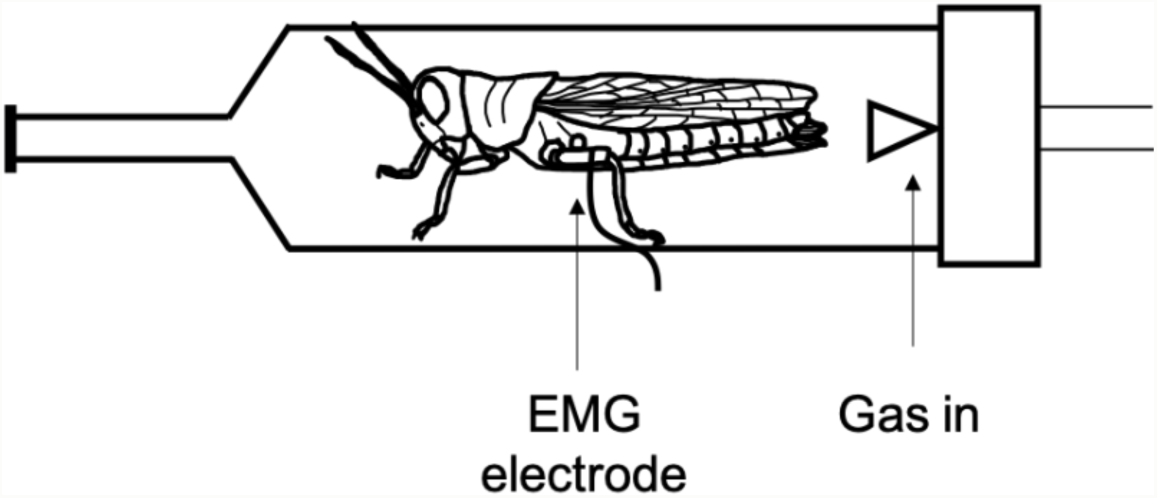
Gas anoxia intact preparation. A schematic diagram of the methods used during gas anoxia experiments. The intact animal is placed within a 50 mL syringe that is fitted with an air supply (gas in) at one end. An EMG electrode was placed in the 3rd abdominal segment, as indicated, to record neural motor patterning and excitability.

### 2.4 Semi-intact preparation

Male and female adult locusts were obtained, appendages and a portion of the pronotum were removed, and a dorsal incision was made along the midline of the animal. Further dissection occurred in a 5 x 2.5 x 2 cm plexiglass chamber with a cork bottom. The dorsal side of the animal was pinned open allowing for removal of the gut, air sacs, and fat bodies and exposure of the ventral nerve cord, including metathoracic ganglion (MTG) and nerve roots (Fig 2A). The abdominal cavity was bathed in standard locust saline containing (in mM): 147 NaCl, 10kCl, 4 CaCl2, 3 NaOH, and 10 HEPES buffer (pH=7.2). A silver ground wire was inserted into the anterior portion of the thorax

**Figure 2.**
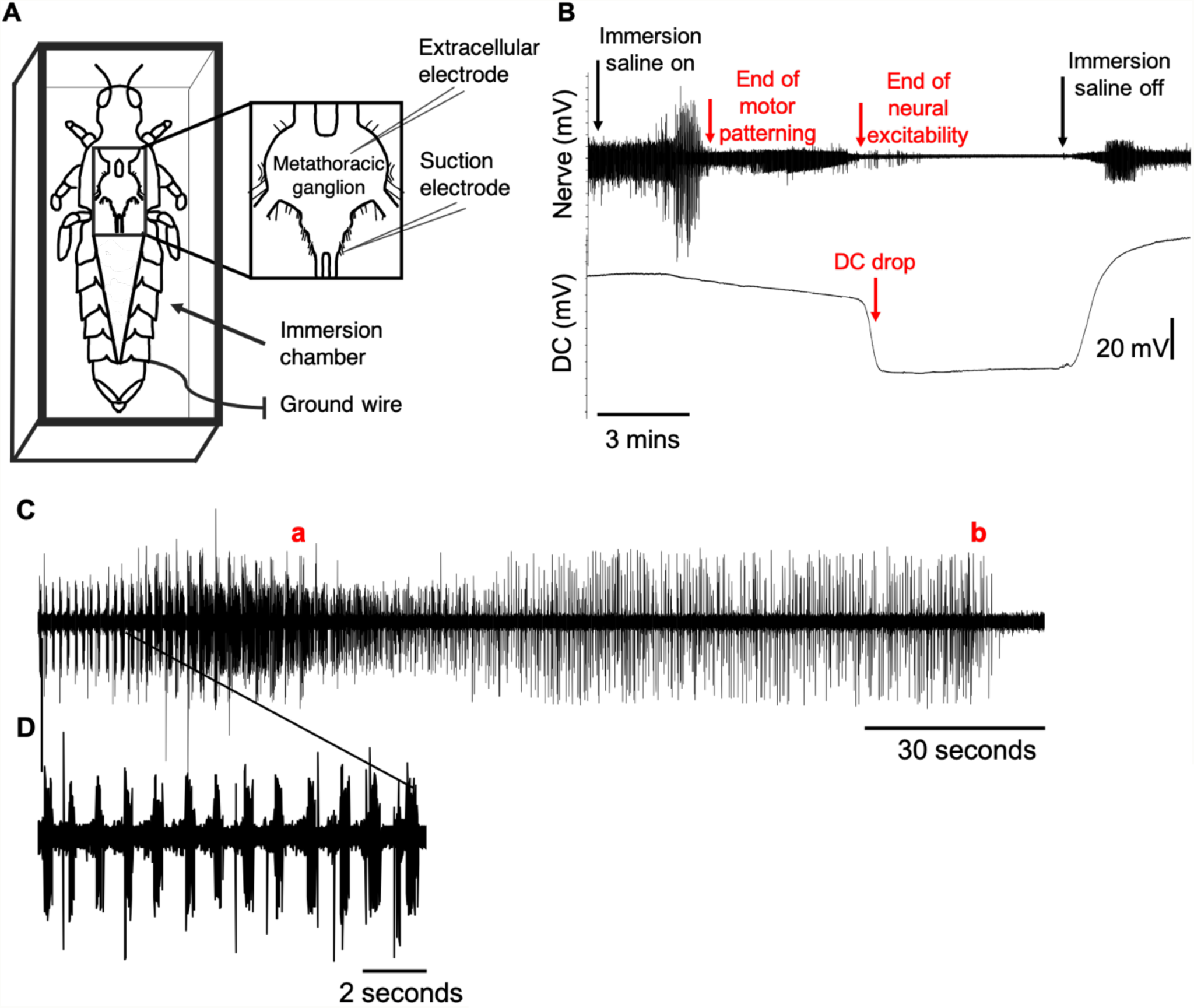
Semi-intact preparation of the locust. (A) Schematic diagram of the dissected animal with extracellular electrode placement in the metathoracic ganglion and a suction electrode attached to a nerve. The immersion chamber is shown, which is filled with saline to induce anoxia. (B) A sample extracellular recording (DC), and nerve recording of a control animal during induction and recovery from anoxic coma via water immersion. (C) A close up of the nerve recording during entry to coma, displaying the end of motor patterning (a) and the end of neural excitability (b). (D) A close up of rhythmic motor patterning.

### 2.5 Electrophysiology

Extracellular microelectrodes were made from filamented capillary tubes (1 mm diameter; World Precision Instruments) pulled to a tip resistance of approximately 5-7 mΩ. Both the electrode and the electrode holder were filled with 3 M KCl and connected to a DC amplifier (model 1600 A-M Systems). Voltage was adjusted to zero with the electrode tip in locust saline, the electrode was then inserted through the sheath of the MTG. Trans-perineurial potential (TPP) was recorded for the entirety of the experiment. Ventilatory motor patterns were recorded from one of the abdominal nerve roots with a suction electrode (World Precision Instruments). The electrode was made from an unfilamented capillary tube 1 mm in diameter (World Precision Instruments) which was pulled to a high-resistance tip before being broken to an appropriate size. The signal was amplified using a differential AC amplifier (model 1700 A-M Systems) and digitized with a DigiData 1322A digitizer (Molecular Devices).

### 2.6 Drug preparation

All chemicals were obtained from Sigma Aldrich Canada and prepared in standard locust saline to concentrations indicated below. Control solutions contain standard locust saline.

### 2.7 Experimental design

#### 2.7.1 Whole animal water immersion

Caffeine treatments consisted of 5 μL, 10 μL, and 15 μL injections of 2.5×10^−2^ M 1, 3, 7- Trimethylxanthine (caffeine), an adenosine receptor antagonist, into the hemocoel at the base of the hindleg. Adenosine treatments consisted of 10 μL injections of 10^−2^ M, 5×10^−2^ M, and 0.1 M 9-β-D-Ribofuranosyladenine (adenosine). Forskolin treatments consisted of 10 μL injections of 10^−2^ M forskolin, a cAMP activator. Control locusts were injected with standard locust saline. During adenosine and forskolin experiments the treatment was injected 20 minutes prior to immersion. During caffeine experiments injection occurred immediately following water emersion to account for half-life and peak concentration, which occur around 15 minutes following injection (National Research Council, 2001).

The time to succumb to coma was measured as the length of time between immersion and the cessation of ventilation and movement. The time to recover was measured as the time between re-exposure to air and the return of ventilation and the ability to stand.

#### 2.7.2 Whole animal gas anoxia

At 10 minutes into the 30-minute coma, the animal was injected with the appropriate treatment and placed immediately back into the syringe. Injections consisted of either 5 μL or 10 μL of 2.5×10^−2^ M caffeine.

The time to succumb to coma was measured as the time between nitrogen gas exposure and the cessation of motor patterning and neural excitability. The time to recovery was measured as the time between re-exposure to air and the time for neural excitability and motor patterning to return.

#### 2.7.3 Semi-intact experiment

The adenosine and caffeine experiments used concentrations of 10^−3^ M and 10^−2^ M for each drug. The NKH 477 and 2’, 3’-dideoxyadenosine (DDA) experiments used drugs at a concentration of 10^−3^ M. NKH-477 was used as a water soluble forskolin derivative to avoid any potential influence of ethanol as a solvent. Following dissection, the animal was bathed in the appropriate treatment for 20 minutes prior to immersion to ensure absorption. The preparation was then flooded with saline to induce anoxia. Immersion saline was left on for 5 minutes and then removed using a large syringe. The time to succumb was measured as the time between immersion and the cessation of neural rhythm production, excitability, and the drop in TPP. The time to recover was measured as the time between immersion saline removal and the return of TPP, neural excitability, and motor patterning (Fig 2B, C).

### 2.8 Statistical analysis

Data were analyzed using SigmaPlot 13 (Systat Software Inc.). One-way and two-way analysis of variance tests were used to determine statistical significance (P < 0.05) between multiple groups of data. Repeated measures tests were used when appropriate. Post hoc tests were used to determine significant differences between individual groups within the data set. The post hoc tests were decided based on the equal variance and normality of the data, measured with Shapiro-Wilk and Levene median tests respectively. Tukey and Holm-Šidák tests were the most commonly used post hoc analyses. Data are reported as mean ± standard deviation or median and interquartile range as appropriate, all data are reported in minutes.

## 3. Results

### 3.1 Whole animal experiments

#### 3.1.1 Adenosine increases time to stand following anoxic coma

Mammalian research demonstrates an increase in adenosine concentrations in the cerebral cortex of mice during spreading depolarization and the activation of ARs lengthen the recovery of synaptic transmission (Lindquist and Shuttleworth, 2012). To determine whether these findings are consistent in insect models we investigated the influence of adenosine on the recovery following anoxic coma. Adenosine had a dose-dependent effect on the recovery of motor control in locusts. The time to succumb was not affected by treatments (One-RM ANOVA P = 0.994). The recovery of ventilation (R_vent_) did not vary across treatments (One-way RM ANOVA P = 0.310) (Fig 3A), however the recovery of standing ability (R_stnd_) was significantly delayed in both adenosine groups (One-way RM ANOVA P < 0.001) (Fig 3B). The time to stand following the return of ventilation also showed a significant difference between groups (One-way RM ANOVA P < 0.001) (Fig 3C).

**Figure 3.**
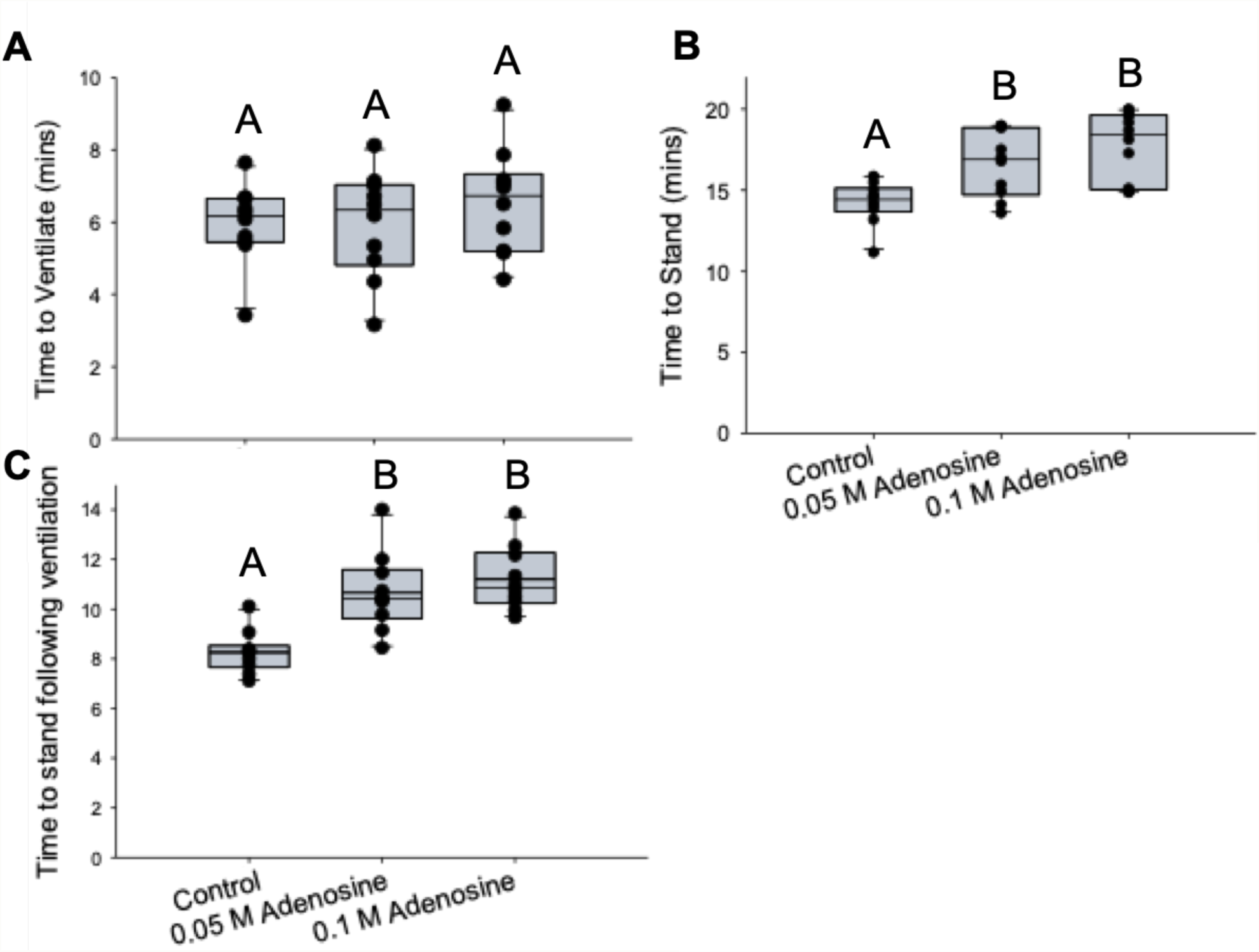
Effect of adenosine on time to recover from anoxic coma via water immersion. (A) The time to recover ventilation (B) Time to stand (C) Time to stand following return of ventilation. All treatments, n=10. Different letters on boxplot indicate P < 0.05.

#### 3.1.2 Caffeine has a dose-dependent, biphasic effect on recovery from anoxic coma

Caffeine acts as an adenosine receptor antagonist, therefore we predicted that increasing doses would decrease the time to recovery, contrasting the effects of adenosine. Caffeine had a biphasic effect on recovery, with the 5 μL dose having the shortest R_vent_ (7.84 ± 1.57 mins) and R_stnd_ (12.46 ± 2.07 mins) (Fig 4A), whereas the 15 μL dose had the longest recovery times (R_vent_11.19 ± 2.02 mins, R_stnd_ 17.11 ± 1.86 mins) (Fig 4B). The control group displayed times intermediate to those from the caffeine groups (R_vent_ 9.63 ± 2.27 mins, R_stnd_ 16.08 ± 3.15 mins). Although there was no statistically significant difference between caffeine 5 μL and control (Holm- Šidák, R_vent_ P = 0.082; Tukey Test, R_stnd_ P = 0.056), the lowest caffeine dose shows a clear decrease in time to recover ventilation and the time to stand.

**Figure 4.**
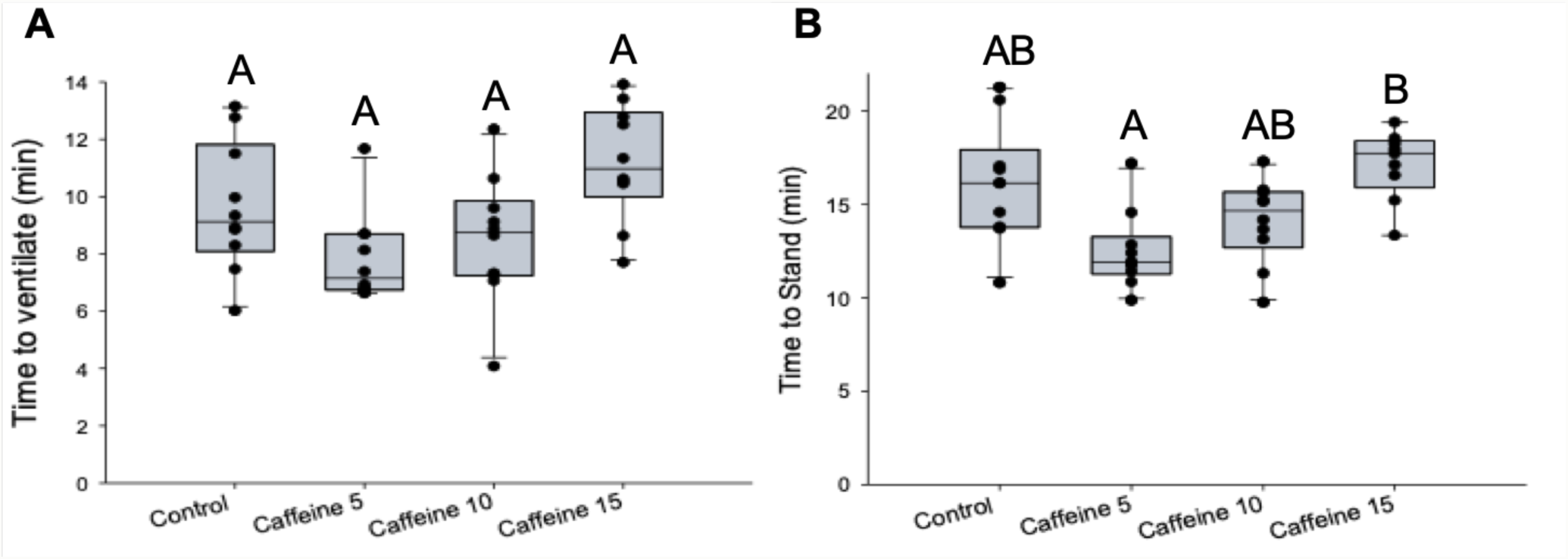
Effect of caffeine on time to recover from anoxic coma via water immersion. (A) The time to recover ventilation (B) Time to stand. All treatments, n=10. Different letters on boxplot indicate P < 0.05.

#### 3.1.3 Caffeine decreases the time to recover neural excitability following gas anoxia

In order to further characterize the effect of caffeine on neural patterning, an electromyographic electrode was used to record ventilatory rhythm during induction to and recovery from anoxic coma. Caffeine treatment groups recovered neural excitability (R_e_) faster than the control group following a 30-minute coma (One-way ANOVA P<0.001) (Fig 5A). Both the 5 μL dose (Holm-Šidák, P < 0.001) and 10 μL dose (Holm-Šidák, P = 0.001), had significantly shorter recovery times than the control (Fig 5B).

**Figure 5.**
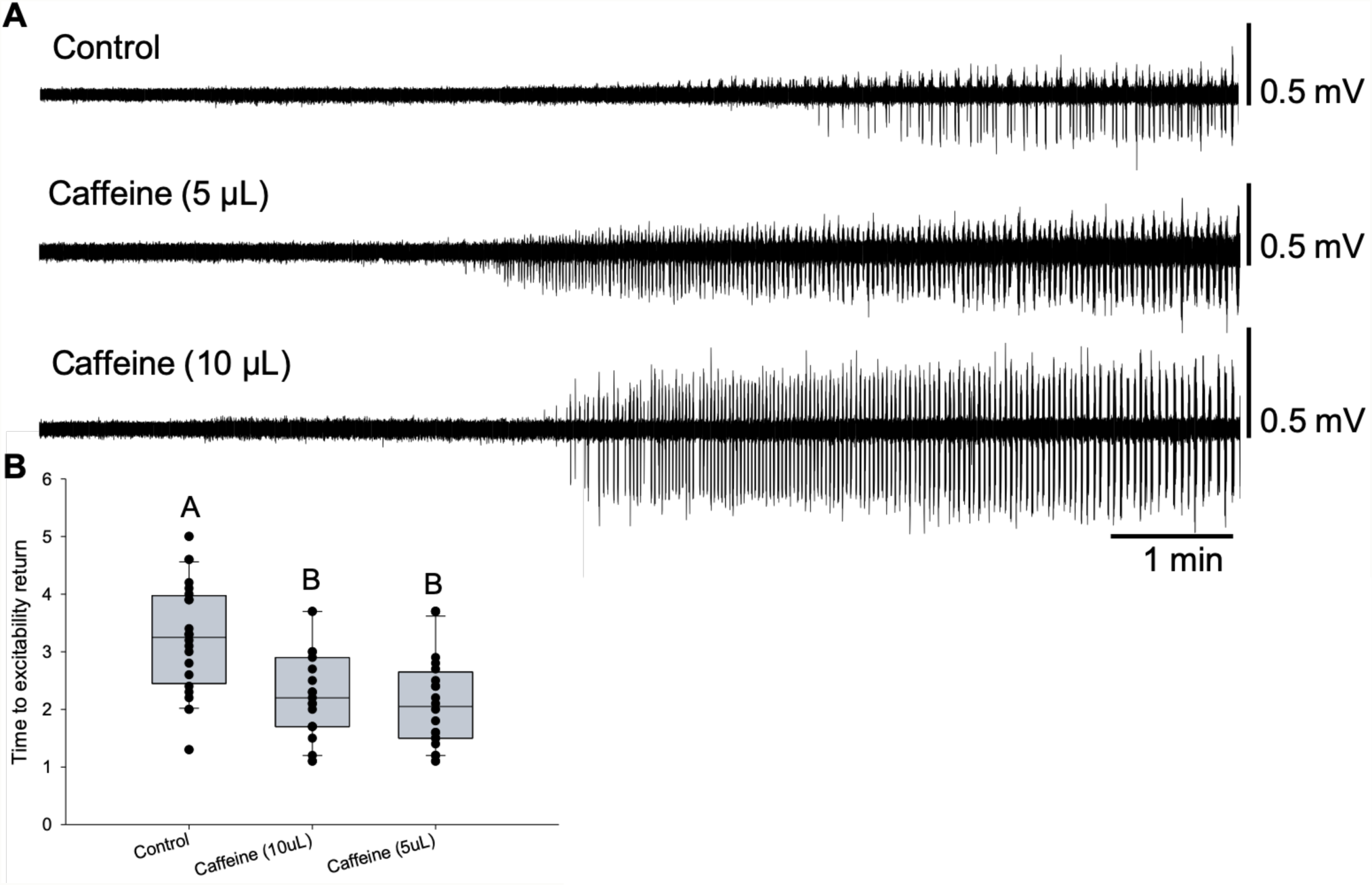
Effect of caffeine on time to recover from anoxic coma via nitrogen gas. (A) Example EMG traces during recovery from coma for each treatment. Each trace begins upon return to air (B) Time to excitability return. All treatments, n=20. Different letters on boxplots indicate P < 0.05.

The return of neural motor patterning (R_mp_) was not significantly different among treatment groups (One-way ANOVA, P = 0.171). R_mp_ was on average faster to recover in caffeine 5 μL (4.56 ± 1.44 mins) than the control group (5.32 ± 1.36 mins), and caffeine 10 μL (4.80 ± 1.27 mins).

Similarly, the return of R_vent_ was quicker on average in caffeine 5 μL (3.9 ± 0.9 mins) and caffeine 10 μL (4.1 ± 1.1 mins), than in controls (4.4 ± 0.8 mins). however, there was no statistical significance (One-way ANOVA P = 0.148).

### 3.2 Semi-intact experiments

#### 3.2.1 Adenosine increases recovery time

Adenosine treatment did not have an effect on the rate of induction to coma (Fig 6A). The R_e_ of animals treated with 1mM adenosine [2.1 (1.15-3.38)] was significantly longer than controls [0.9 (0.45-1.45)] (Dunn’s Method, P = 0.012) (Fig 6B). R_mp_ was not significantly influenced by adenosine treatment (Fig 6C). The timing of TPP was not affected by the adenosine treatments.

**Figure 6.**
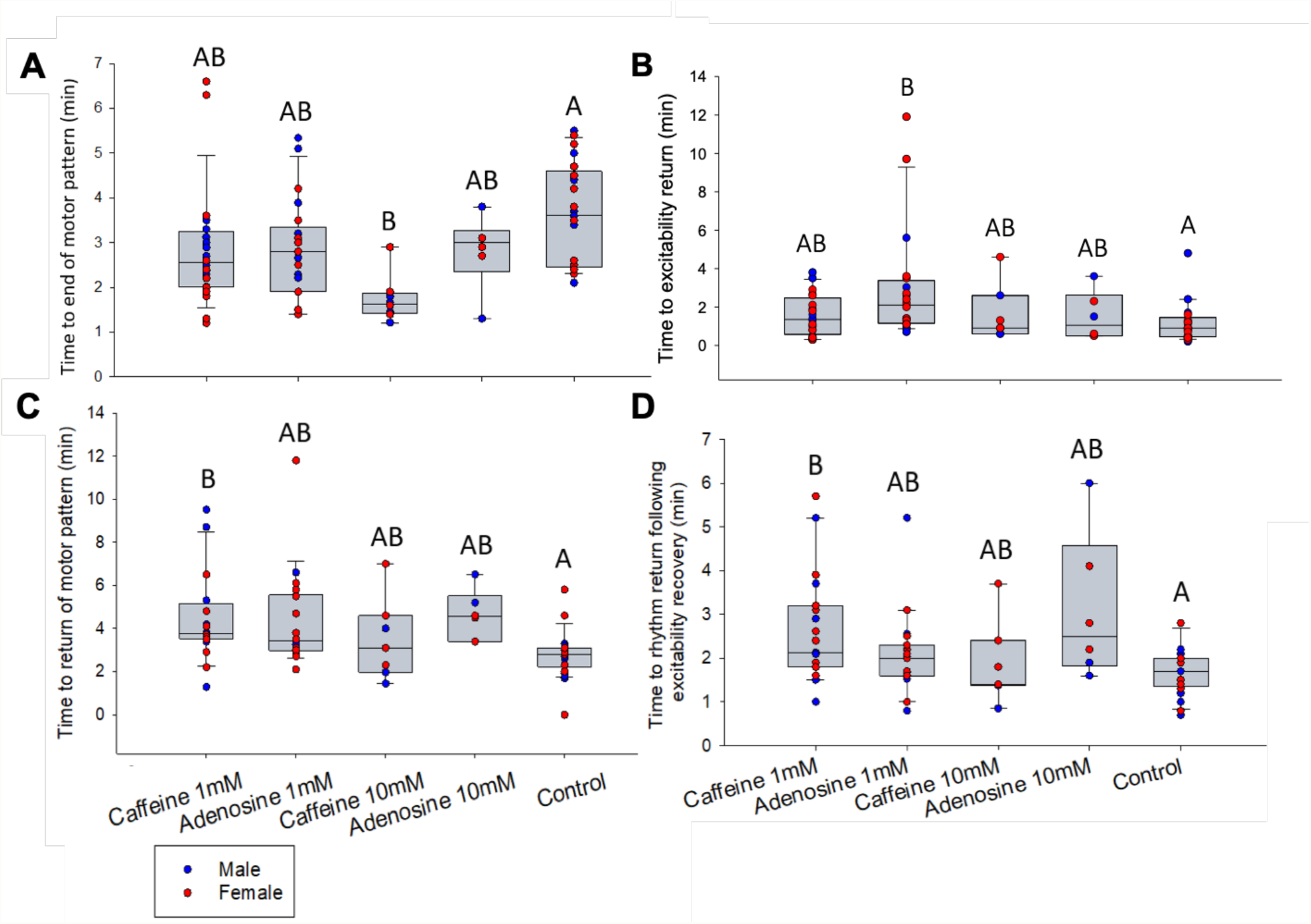
Effect of varying concentrations of adenosine and caffeine on time to succumb and time to recover from anoxic coma via water immersion. (A) The time to lose motor patterning during coma induction. (B) The time to return of neural excitability. (C) The time to return of motor patterning. (C) The time to return of motor patterning following return of excitability. Caffeine 1mM, adenosine 1 mM, and control n=10, per sex; Caffeine 10 mM and adenosine 10 mM, n=3, per sex. Different letters on boxplots indicate P < 0.05. Details in text.

#### 3.2.2 Caffeine affects coma induction and recovery rates

Compared to controls, animals treated with 10^−2^ M caffeine lost neural excitability sooner (Two-Way ANOVA, P = 0.043) and lost neural motor patterning (F_mp_) sooner (Two-Way ANOVA P = 0.001) during coma induction (Fig 6A).

Animals treated with 10^−3^ M caffeine had a significantly longer R_mp_ (4.89 ± 0.50 mins) compared to controls (2.76 ± 0.49 mins) (Holm-Šidák, P = 0.038) (Fig 6C). The return of motor patterning following the return of neural excitability (R_mp-e_) was significantly longer in 10^−3^ M caffeine treatments (2.93 ± 0.27) compared to the controls (1.69 ± 0.27) (Holm-Šidák, P = 0.019) (Fig 6D). The timing of R_e_ and TPP were not affected by the caffeine treatments.

#### 3.2.3 Adenylyl cyclase activation increases recovery time in intact animals

Preliminary research showed that forskolin, an adenylyl cyclase activator, increased R_stnd_ compared to controls (One-Way ANOVA P = 0.034). NKH-477 was used as a forskolin derivative, and DDA was used as an adenylyl cyclase inhibitor. NKH-477 had a longer R_mp_ (5.29 ± 0.52 mins) compared to controls (2.76 ± 0.35 mins) (Holm-Šidák, P = 0.001), however DDA did not show a significant difference (Fig 7A).

**Figure 7.**
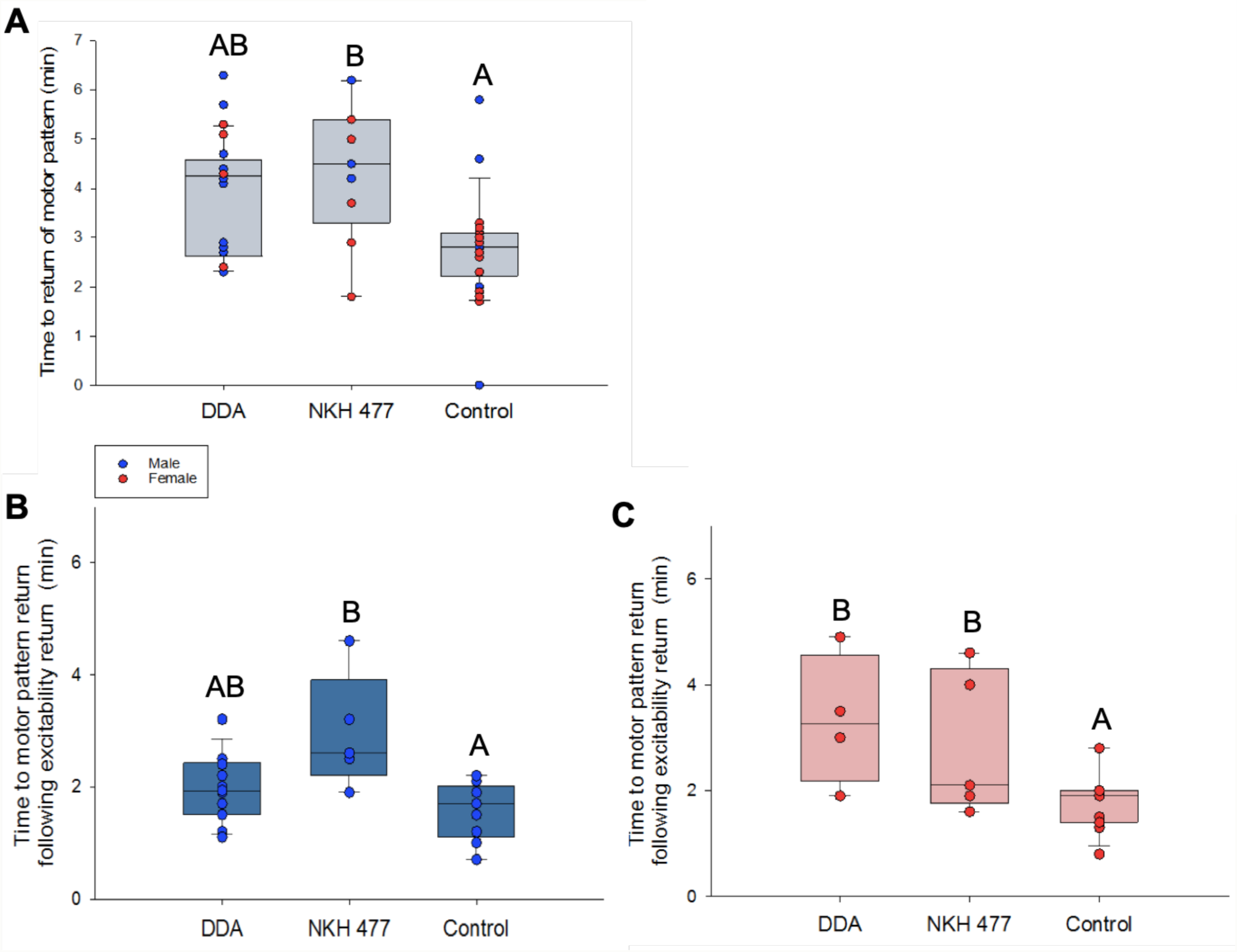
Effect of DDA and NKH-477 on recovery from anoxic coma via water immersion (A) The time to motor pattern recovery (B) The time to motor pattern return following the return of excitability in males (C) The time to motor pattern return following the return of excitability in females. DDA male n=19, female n=5; NKH-477 n=5, per sex. Control n=10 per sex. Different letters on boxplots indicate P < 0.05. Details in text.

R_mp-e_ was influenced by both treatments. However, there was a significant difference between sexes (Holm-Šidák, P = 0.002). The DDA treatment increased R_mp-e_ compared to controls in females only (Holm-Šidák, P = 0.007). The NKH-477 treatment increased R_mp-e_ compared to controls in both males (Holm-Šidák, P = 0.009), and females (Holm-Šidák, P = 0.041) (Fig 7B).

## 4. Discussion

The mechanisms involved in SD propagation and recovery are not completely known and we investigated the role of adenosine in locusts. We found that manipulation of the adenosine receptor influences the timing of anoxic coma induction and recovery. Administration of adenosine as a receptor agonist delayed recovery following coma in both intact and semi-intact preparations. However, treatment with caffeine (AR antagonist), had mixed results. There was a dose-dependent effect in intact locusts; lower concentrations shortened, and higher concentrations lengthened recovery times. In semi-intact preparations caffeine hastened induction to coma and delayed recovery. An activator of cAMP and an inhibitor both delayed recovery.

### 4.1 Adenosine

Adenosine has been linked to the prolonged depression of synaptic transmission following the restoration of ion gradients in mammalian models (Lindquist and Shuttleworth, 2012). The build-up of extracellular adenosine contributes to the delay in recovery of synaptic transmission during SD recovery. Adenosine production is dependent on metabolic state; under physiological conditions, production primarily occurs intracellularly, however, when energy levels are low extracellular production is increased (Deussen et al; 2000, 1999). The primary mechanism for extracellular adenosine production is dephosphorylation of its precursor molecules: ATP, ADP, and AMP. Vertebrates have four adenosine receptors (ARs), all with different downstream pathways. Due to the lack of research surrounding invertebrate AR function, it was unknown whether activation would result in similar effects as found in mammals. We show that adenosine does lengthen the time of neural recovery following anoxic coma.

Our initial results demonstrate a clear effect of adenosine on functional recovery. Only males were used during whole-animal experiments to establish a clear pharmacological effect, independent of sex. The time to stand in an intact animal depends on the recovery of synaptic transmission and motor pattern generation. Our results agree with predictions, showing that increased adenosine delays the time to stand, an effect that is exacerbated with an increased concentration. This effect was not present in the time to succumb to coma or the time to recover ventilation. This suggests that adenosine has differential roles and that some mechanisms are independent of its effects. Entry to coma is dependent on the suppression of synaptic activity and disruption of ionic gradients (Somjen, 2001). This process is associated with surges of extracellular K^+^, and a decrease in extracellular sodium, chloride and calcium (Pietrobon and Moskowitz, 2014). AR activation during this time may not have a large effect due to the significant consequences of ion channel disruption. The ventilatory central pattern generator (vCPG) is located within the MTG, allowing ventilation to act as a qualitative measure of neuromuscular function during entry and recovery to coma (Rodgers 2007). However, the time to recover ventilation was not greatly affected by AR activation. This may be a consequence of the mode of anoxia. Compared to gas or chemical anoxia, water immersion requires additional recovery time to clear excess water from around the spiracles. This process may influence the timing of ventilatory recovery, even in the presence of pharmacological agents.

A semi-intact preparation allowed for the measurement of trans-perineurial membrane potential and the negative DC shift associated with SD. An additional recording from a ventilatory nerve assessed changes in neural excitability and motor pattern generation. Both male and female locusts were used for semi-intact experiments in order to determine whether sex has an influence. A bath application of adenosine influenced the return of neural excitability. Excitability return was delayed in treatment groups compared to controls. This finding aligns with our predictions that adenosine lengthens recovery times. Although females seemed to be affected more by treatment than males, there was no difference due to sex.

The return of motor patterning was not influenced by adenosine treatments. This was not predicted. The semi-intact preparation is more invasive, and this may impact motor pattern generation. The insect central nervous system is covered by a thin muscular septum known as the ventral diaphragm (Richards, 1963). This diaphragm is composed of a single layer of muscle fibres, which play an important role in ventilation as their electrical activity is coupled to that of the main inspiratory muscles (Peters, 1977). In semi-intact preparations this diaphragm is removed, exposing the connective-tissue sheath covering the MTG. Due to its rhythmical activity and connection with ventilatory movement, the removal of the diaphragm may impact the recovery of motor patterning. Although there was an influence of adenosine on the return of excitability, motor pattern generation requires coordinated signalling that may be more greatly affected by the disruption of dissection than that of adenosine application.

### 4.2 Caffeine

Caffeine acts as a non-selective AR antagonist and it elicits similar effects in invertebrates as in mammals (Mustard, 2014; Wu et al, 2009). Although AR antagonism is the primary mechanism of action, caffeine also acts to inhibit phosphodiesterase, and to mobilize intracellular calcium (Nehlig et al, 1992). Our results show that caffeine has a biphasic effect, with low dosages speeding up time to recovery and high dosages delaying recovery. In mammalian studies this is attributed to the binding affinity of ARs (Nehlig et al, 1992; Pedata et al, 1984). Similar to adenosine, at low concentrations caffeine will more readily bind to high affinity ARs. When caffeine concentrations are high, low affinity ARs will also be bound. Although only one AR has been identified in invertebrates, a similar effect may be occurring through interactions with other receptors. As a result of the wide scope of caffeine’s actions, high doses may be interacting with other molecules and causing an inhibitory effect on the nervous system.

Caffeine administration at a low dose decreased the average time to recover excitability and motor patterning following exposure to gas anoxia. These findings were not as strong as previous results. The placement of the EMG electrode introduced some variability. The electrode was placed in the same approximate location on each animal; however, the depth of the electrode tip could have varied up to 3 mm. Due to this variability, we may have recorded from different muscles, causing inconsistencies.

Drug preparation took into account the volume of haemolymph in each animal, which results in a dilution of the administered solution. Due to variation in body size, haemolymph volume is 142 – 369 μL for adult male locusts (Loughton and Tobe, 1969; Ayali and Penner, 1992). Our concentrations were prepared based on an average haemolymph volume of 200 μL. This variation in haemolymph volume may account for irregularities within whole animal experiments.

In semi-intact preparations caffeine increased the time to coma induction and delayed the time to recovery. These results indicate that caffeine contributed to the ionic disturbance associated with the onset of anoxic coma. Both rhythm and excitability were lost sooner in high concentration caffeine treatments (10^−2^ M). Although it is evident that a loss of ion gradients is critical to the initiation of SD, it is still unclear what mechanisms are behind this initiation and propagation. Both mammalian and invertebrate research shows that SD occurs first as neuronal silencing followed by membrane depolarization (Rodgers et al, 2010; Muller and Somjen, 2000). Our findings show that caffeine influences neural excitability and rhythm generation, however, it has no effect on the timing of the DC shift. This suggests that caffeine has an effect on the inactivation of neurons and not membrane depolarization.

Lower concentration caffeine treatments (10^−3^ M) delayed the recovery of neural rhythms but not that of excitability. Contrary to our predictions, this finding indicates that caffeine acted to slow functional recovery at the synaptic level. These findings are surprising given our previous research on intact animals. The semi-intact preparation is more invasive and results in increased damage to the animal, further exposing the ganglia and nervous system to each treatment. Due to this increased exposure, the low concentration caffeine had greater access to the ganglia and therefore increased potential for binding to receptors. This may have been the cause for the delay in recovery, similar to our preliminary findings. Further research will investigate the effects of lower concentrations in a semi-intact preparation to confirm this finding.

### 4.3. Adenylyl cyclase

AR1 activation results in the inactivation of adenylyl cyclase, the enzyme responsible for cAMP production. cAMP is involved in protein kinase A (PKA) activation. This pathway has been shown to reduce recovery times following hyperthermic coma, mediated by octopamine (Armstrong et al 2006). We explored whether adenylyl cyclase activation and inhibition would produce similar results as the AR manipulations. Preliminary findings with intact animals showed that forskolin, an adenylyl cyclase activator, delayed the time to stand following coma. Similarly, a water-soluble forskolin derivative, NKH-477, increased the time to recover neural rhythm in a semi-intact preparation. These findings were surprising, as previous research had demonstrated that an increase in cAMP improved recovery times following hyperthermic coma (Armstrong et al, 2006). The preliminary research used a forskolin concentration higher than that of previous studies, to ensure drug permeability in an intact preparation. This concentration may have been too high, resulting in a delay in the time to stand. A similar effect may have arisen in the semi-intact preparations. Perhaps this increased access resulted in an overwhelming effect similar to that of high concentrations of caffeine. DDA was used as an adenylyl cyclase inhibitor, which delayed the return of rhythm following the return of excitability in females. This variation between sexes may be a result of differences in behaviour. When reared in a crowded colony, adult male locusts must compete for female mates. This is a behaviour that requires additional neural activity and hormonal control. This is supported by previous results that show an increase in the time to ventilate in males during following sexual maturity (Robertson et al, 2019). This finding agrees with our predication that decreased cAMP will delay recovery.

In conclusion, our results demonstrate that increased adenosine does affect the recovery of motor patterning in invertebrates. Motor pattern recovery is reliant on functional synapses suggesting that increased adenosine delays synaptic recovery. Adenosine administration resulted in the delay of functional recovery. This finding was shown in both intact and semi-intact preparations. We also found that sex affects the recovery of neural excitability. Caffeine as an AR antagonist showed contrasting results; due to the wide mechanistic scope of this drug, additional research requires greater variation in treatment concentrations. These findings indicate that increased adenosine concentrations have a similar effect on the recovery of synaptic transmission as was found in mice. The effects of AR manipulation may be mediated by the cAMP pathway, as similar findings are shown through adenylyl cyclase activation and inhibition. A clear finding from our results is that the timing of the trans-perineurial potential drop and recovery was stable and not influenced by our pharmacological treatments. This indicates a clear distinction between the mechanisms responsible for recovery of membrane potential and the effects of adenosine on synaptic recovery.

## Abbreviations

AR: Adenosine receptor
F_mp_: Failure of motor patterning
MTG: Metathoracic ganglion
R_e_: The recovery of neural excitability
R_mp_: The recovery of motor patterning
R_mp-e_: The recovery of motor patterning following the recovery of excitability
R_stnd_: The recovery of standing ability
R_vent_: The recovery of ventilation
SD: Spreading depolarization
TPP: Trans-perineurial potential

## References

Andretic, R., Kim, Y. C., Jones, F. S., Han, K. A., & Greenspan, R. J. (2008). Drosophila D1 dopamine receptor mediates caffeine-induced arousal. Proceedings of the National Academy of Sciences, 105, 20392–20397.

Armstrong, G. A., Shoemaker, K. L., Money, T. G., & Robertson, R. M. (2006). Octopamine mediates thermal preconditioning of the locust ventilatory central pattern generator via a cAMP/protein kinase A signaling pathway. Journal of Neuroscience, 26, 12118–12126.

Ayali, A., & Pener, M. P. (1992). Density-dependent phase polymorphism affects response to adipokinetic hormone in Locusta. Comparative Biochemistry and Physiology Part A: Physiology, 101, 549–552.

Borea, P. A., Gessi, S., Merighi, S., Vincenzi, F., & Varani, K. (2018). Pharmacology of adenosine receptors: the state of the art. Physiological reviews, 98, 1591–1625.

Deussen, A. (2000). Metabolic flux rates of adenosine in the heart. Naunyn-Schmiedeberg’s archives of pharmacology, 362, 351–363.

Deussen, A., Stappert, M., Schäfer, S., & Kelm, M. (1999). Quantification of extracellular and intracellular adenosine production: understanding the transmembranous concentration gradient. Circulation, 99, 2041–2047.

Dolezelova, E., Nothacker, H. P., Civelli, O., Bryant, P. J., & Zurovec, M. (2007). A Drosophila adenosine receptor activates cAMP and calcium signaling. Insect Biochemistry and Molecular Biology, 37, 318–329.

Fredholm, Bertil B., Adriaan P. IJzerman, Kenneth A. Jacobson, Karl-Norbert Klotz, and Joel Linden. (2001) International Union of Pharmacology. XXV. Nomenclature and classification of adenosine receptors. Pharmacological Reviews, 53, 527–552.

Hendricks, J. C., Finn, S. M., Panckeri, K. A., Chavkin, J., Williams, J. A., Sehgal, A., & Pack, A. I. (2000). Rest in Drosophila is a sleep-like state. Neuron, 25, 129–138.

Kalinowski, R. R., Jaffe, L. A., Foltz. K. R., Giusti AF (2003) A receptor linked to a Gi-family G-protein functions in initiating oocyte maturation in starfish but not frogs. Developmental Biology, 253, 139–149

Lindquist, B. E., & Shuttleworth, C. W. (2012). Adenosine receptor activation is responsible for prolonged depression of synaptic transmission after spreading depolarization in brain slices. Neuroscience, 223, 365–376.

Lindquist, B. E., & Shuttleworth, C. W. (2017). Evidence that adenosine contributes to Leao’s spreading depression in vivo. Journal of Cerebral Blood Flow & Metabolism, 37, 1656–1669.

Loughton, B. G., & Tobe, S. S. (1969). Blood volume in the African migratory locust. Canadian Journal of Zoology, 47, 1333–1336.

Müller, M., & Somjen, G. G. (2000). Na+ and K+ concentrations, extra-and intracellular voltages, and the effect of TTX in hypoxic rat hippocampal slices. Journal of neurophysiology, 83, 735–745.

Mustard, J. A. (2014). The buzz on caffeine in invertebrates: effects on behavior and molecular mechanisms. Cellular and molecular life sciences, 71, 1375–1382.

National Research Council. (2001). Caffeine for the sustainment of mental task performance: Formulations for military operations. Washington, DC: National Academy Press, 6, 104–168.

Nehlig, A., Daval, J. L., & Debry, G. (1992). Caffeine and the central nervous system: mechanisms of action, biochemical, metabolic and psychostimulant effects. Brain Research Reviews, 17, 139–170.

Pedata, F., Pepeu, G., & Spignoli, G. (1984). Biphasic effect of methylxanthines on acetylcholine release from electrically-stimulated brain slices. British journal of pharmacology, 83, 69.

Peters, M. (1977). Innervation of the ventral diaphragm of the locust (Locusta migratoria). Journal of Experimental Biology, 69, 23–32.

Pietrobon, D., & Moskowitz, M. A. (2014). Chaos and commotion in the wake of cortical spreading depression and spreading depolarizations. Nature Reviews Neuroscience, 15, 379–393.

Richards, A. G. (1963). The ventral diaphragm of insects. Journal of Morphology, 113, 17–47.

Robertson, R. M., Cease, A. J., & Simpson, S. J. (2019). Anoxia tolerance of the adult Australian Plague Locust (Chortoicetes terminifera). Comparative Biochemistry and Physiology Part A: Molecular & Integrative Physiology, 229, 81–92.

Rodgers, C. I., Armstrong, G. A., & Robertson, R. M. (2010). Coma in response to environmental stress in the locust: a model for cortical spreading depression. Journal of insect physiology, 56, 980–990.

Rodgers, C. I., Armstrong, G. A., Shoemaker, K. L., LaBrie, J. D., Moyes, C. D., & Robertson, R. M. (2007). Stress preconditioning of spreading depression in the locust CNS. PloS one, 2, e1366.

Rodgers, K. I., LaBrie, J. D., & Robertson, R. M. (2009). K+ homeostasis and central pattern generation in the metathoracic ganglion of the locust. Journal of insect physiology, 55, 599–607.

Somjen, G. G. (2001). Mechanisms of spreading depression and hypoxic spreading depression-like depolarization. Physiological reviews, 81, 1065–1096.

Spong, K. E., Mazzetti, T. R., & Robertson, R. M. (2016). Activity dependence of spreading depression in the locust CNS. Journal of Experimental Biology, 219, 626–630.

Wu, M. N., Ho, K., Crocker, A., Yue, Z., Koh, K., & Sehgal, A. (2009). The effects of caffeine on sleep in Drosophila require PKA activity, but not the adenosine receptor. Journal of Neuroscience, 29, 11029–11037.

